# A Computational Approach to Interpreting the Embedding Space of Dimension Reduction

**DOI:** 10.1101/2024.06.23.600292

**Authors:** Bingyuan Zhang, Kohei Uno, Hayata Kodama, Koichi Himori, Yusuke Matsui

## Abstract

Nonlinear dimension reduction methods are widely applied in studies analyzing gene and protein expression, by revealing patterns of discrete groups and continuous orders in high-dimensional data. However, the tools are limited to understanding the obtained embedding structures of biological mechanisms, hindering the full exploitation of data. Here, we propose a novel framework to interpret embedding systematically by identifying and mapping associated biological functions. The method performs statistical tests and visualizes significantly enriched functions essential for the organization of the embedding structure, by applying it to the embedding results of two datasets: the Genotype Tissue Expression dataset and a *Caenorhabditis elegans* embryogenesis dataset, one capturing distinct cluster structures and the other capturing continuous developmental trajectories. We identified the associated functions for interpreting the two embeddings and confirmed it as a useful explainable AI tool in exploratory data analysis by providing annotations to the embedding space.

## Introduction

Dimension reduction (DR) methods are powerful tools for exploratory data analysis in bulk and single-cell transcriptomics, as well as other high-dimensional omics datasets. These methods transform data from a high-dimensional space to a lower-dimensional embedding space, enabling easier visualization and providing an intuitive understanding of the data. The results of dimension reduction are influenced by the chosen method, which can be categorized as either linear or nonlinear. Linear methods, such as Principal Component Analysis (PCA) and Multidimensional Scaling (MDS), are favored for their interpretability. However, these methods often fail to reveal hidden structures in complex high-dimensional data, as observed in the first two principal components^1^. Contrastingly, nonlinear DR methods, also known as manifold learning methods, such as t-distributed stochastic neighbor embedding (t-SNE)^2^ and uniform manifold approximation and projection (UMAP)^3^, have gained popularity. These nonlinear methods aim to preserve the local neighborhood structures when embedding the sample points, enabling the identification of hidden clusters and capturing developmental biological processes in 2D/3D spaces^2–6^. However, the embedding results of nonlinear DR methods are sensitive to hyperparameters, and interpreting the transformation from high-dimensional to low-dimensional space is challenging. Several studies have guided applying these methods to practical data analysis^1,7,8^. Recent studies have focused on exploring ways to better preserve global and local data structures through algorithmic advancements^9–11^. Despite these advancements, understanding the embedding results from a biological perspective remains difficult, making the embedding structure of a high-dimensional dataset a black box for application users.

The lack of interpretability in nonlinear models has fast-tracked the development of the field of explainable AI (XAI)^12^. Various model-agnostic methods, such as LIME^13^ and SHAP^14^ have been proposed to enhance the transparency of nonlinear models such as neural networks. These post-hoc methods elucidate the prediction process of a black-box model by identifying important features for individual instances. However, current XAI methods in biological applications mainly focus on predictive models within a supervised learning framework, assuming the availability of label information^15^. Few XAI methods have been designed for dimension reduction, which belong to the unsupervised learning framework, to explore the biological insights of the embedding space. A recent study adapted the LIME framework to identify the important features for t-SNE embedding results^16^, but it did not address the links between the embedding results and related biological mechanisms and functions. Understanding the embedding results of nonlinear DR methods remains a challenge in machine learning^12^. In biological research, there is a need for an XAI tool that can elucidate the biological mechanisms underlying data analysis results. This study aims to interpret embedding results in terms of the associated biological functions. Despite the widespread application of DR methods to transcriptome and proteomic datasets, mapping related biological functions to the embedding space remains an open problem.

We developed a computational framework called Post-hoc Interpretation of Embedding (PIE) for exploratory data analysis. PIE offers a systematic post-hoc analysis of embeddings through functional annotation, identifying and visualizing the biological functions associated with the embedding structure. The process begins by mapping genes to the embedding space based on the original expression matrix and the embedding coordinates of the samples. These mapped genes are then consolidated to enable biological interpretation. By mapping related functions to an embedding space, PIE offers a novel perspective for exploring embedding structures in a visualizable and comprehensible manner.

Numerical experiments demonstrate that PIE effectively identifies functions that adequately interpret the embeddings. Current literature often focuses on applying DR to biological data to capture distinct clusters (e.g., phenotypes) and continuous developmental processes (e.g., cell differentiation trajectories). To validate our method, we addressed two biological problems using datasets: the Genotype Tissue Expression (GTEx) dataset^17^ and the *Caenorhabditis elegans* embryogenesis dataset^18^. The GTEx dataset, a bulk RNA-seq dataset, has embeddings that capture separated clusters of different tissue types. The *C. elegans* dataset, a single-cell RNA-seq dataset, has embeddings that capture the developmental trajectories of differentiated cell type lineages from the early embryo stage. We used Gene Ontology Biological Process (GOBP) terms^19^ to map and interpret these embeddings. Our comprehensive experiments illustrate the effectiveness of PIE in interpreting both distinct clusters and continuous developmental trajectories, showcasing its utility in providing functional annotations that make the embeddings more interpretable.

## Results

### Overview of PIE

An overview of the proposed computational pipeline is shown in Figure 1. PIE uses three inputs: (A1) a low-dimensional embedding matrix, (A2) an expression matrix, and (A3) functional gene sets of interest (**Figure 1A**). The embedding matrix contains the 2D/3D coordinates of samples obtained using the nonlinear DR method, with UMAP^3^ as the default choice for dimension reduction. The expression matrix can be derived from bulk RNA-seq or other variations such as single-cell RNA-seq or proteomic expression matrices. Functional gene sets can be user-defined or sourced from public databases. For this study, we used the Gene Ontology (GO) knowledgebase^19,20^, which offers extensive information on gene sets related to biological functions. PIE proceeds through the following steps to achieve the annotation of an embedding.

**Figure 1.**
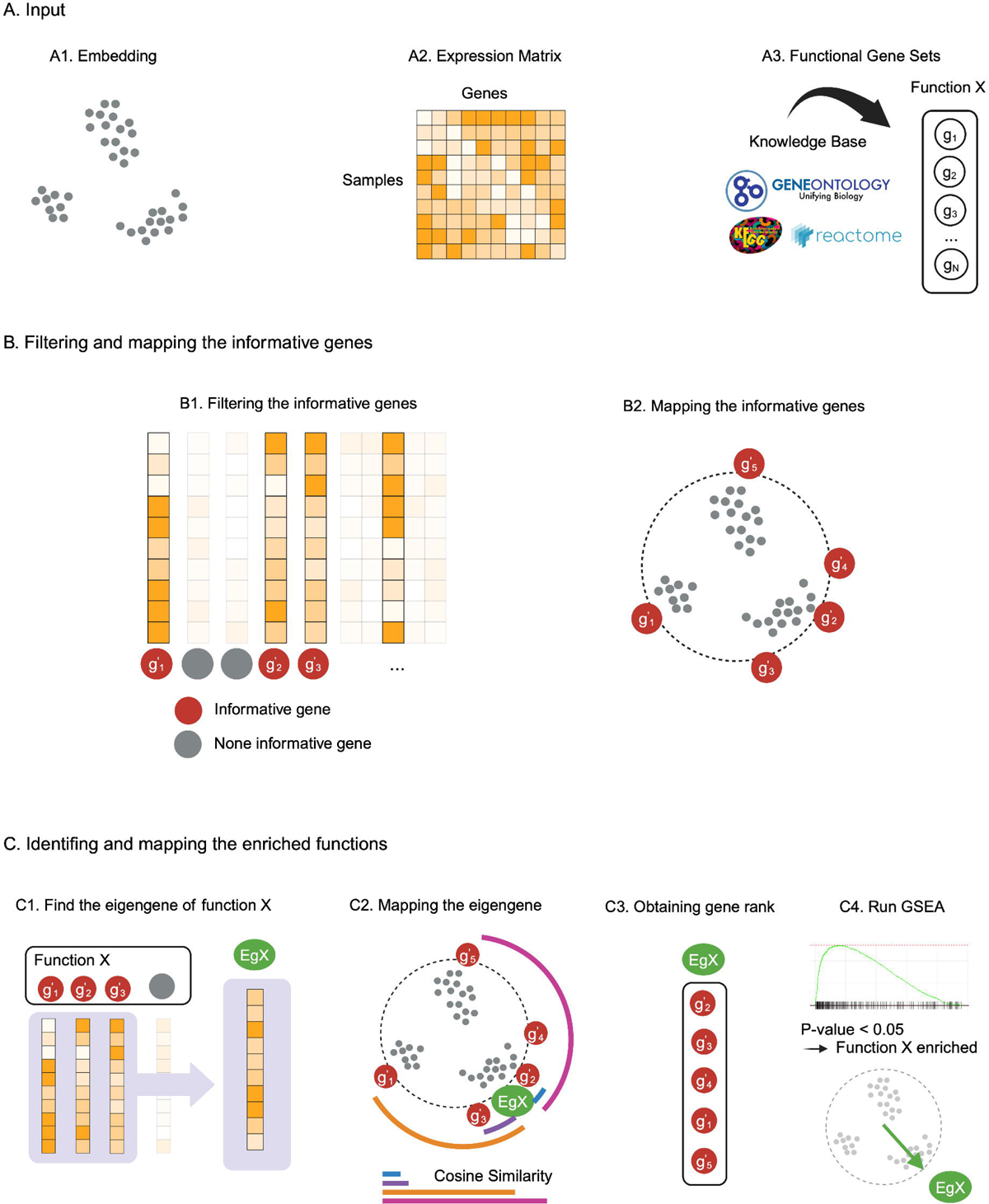
Overview of PIE. A. The three inputs of PIE are (A1) a low-dimensional embedding, (A2) an expression matrix and (A3) functional gene sets. Low-dimensional embedding is obtained by reducing the dimensions of the expression matrix using a nonlinear DR method such as UMAP. Functional gene sets consist of gene lists that are known to be associated with biological functions, which can be either user-defined or sourced from public knowledge bases, such as Gene Ontology^19^, KEGG^39^ and Reactome^40^. In the following analysis, we use Gene Ontology knowledge as an example. B. Filtering and mapping of informative genes to the embedding space. Genes that showed higher consistency with the embedding structure were termed informative and selected for subsequent analysis. The selected informative genes were mapped to the embedding space using projection pursuit. C. Identifying and mapping enriched functions to embedding space. For a functional gene set, we focused on the informative genes in the set and calculated eigengenes to characterize the expression patterns of the functional gene set. Next, we mapped the eigengenes like the mapping of single genes using projection pursuit. Based on the mapped coordinates of the eigengene, we calculated the cosine similarities between the eigengene and other mapped genes, which allowed us to define a gene rank based on similarity measures. Using the obtained gene rank, we applied the Gene Set Enrichment Analysis (GSEA) test^25^ to identify whether function X was enriched towards the mapped direction. If function X was significantly enriched, an arrow pointing to its mapped coordinates was used to visualize its enriched direction.

Initially, PIE filters informative gene vectors (**Figure 1B**). In the literature, gene vectors that exhibit higher consistency with the embedding are termed “informative.” This step identifies genes that best preserve information regarding the embedding structure of a dataset. PIE adopts SCMarker^21^, a filter designed for single-cell RNA-seq datasets but also applicable to bulk RNA-seq datasets, to identify these informative genes, which serve as discriminative markers for capturing the data structure (**Figure 1B, B1**). Next, PIE maps the selected genes to the embedding space using projection pursuit^22,23^ (**Figure 1B, B2**). Projection pursuit determines a linear projection that maximizes the association between the embedding coordinates and each gene vector. The mapped coordinates of genes are similar to the loading vectors used to visualize a PCA biplot. These mapped coordinates are denoted by weighting vectors in projection pursuit, representing the associated direction of genes in the embedding space (**see method details**). When performing projection pursuit, the maximized correlation indicates the consistency between the embedding and genes from a global perspective. Therefore, PIE applies an additional filter to focus on gene vectors with higher maximized association measures. After filtering, the normalized weighting vectors represent the corresponding genes on a unit circle/sphere in the 2D/3D embedding space.

Finally, PIE identifies and visualizes biological functions within the embedding space (**Figure 1C**). For each functional gene set, PIE identifies overlapping informative genes (**Figure 1C, C1**) and calculates the eigengene^24^ by integrating the expression patterns of these overlapping genes (**Figure 1C, C2**). The eigengenes are then mapped to the embedding space. Since both mapped genes and eigengenes reside on a unit circle, their cosine similarities can be computed. A higher cosine similarity indicates a strong resemblance between the mapped coordinates of the eigengene and gene. PIE ranks the mapped informative genes for each functional gene set based on these similarities (**Figure 1C, C3**). Using these ranks, PIE performs a Gene Set Enrichment Analysis (GSEA) test^25^ to identify if the functional gene sets are significantly enriched (p < 0.05) (**Figure 1C, C4**). After statistical testing, PIE provides two visualizations for enriched functions: an arrow pointing in the enriched direction of the corresponding function and a sample-wise functional enrichment plot showing the relative expression level of a gene set (**see method details**). Thus, PIE provides an annotation of the embedding.

### PIE interprets the tissue-specificity in the GTEx embedding

For the first problem, we used the Genotype Tissue Expression (GTEx) dataset^17^, a bulk RNA-Seq dataset. The 2D GTEx embedding, obtained using UMAP, revealed distinct tissue clusters (**Figure 2A**). Applying PIE to GTEx embedding demonstrated the following key points: (1) PIE identified the associated functions and mapped their enriched directions in the embedding space. Moreover, the enriched directions of functions characterized the functional differences between tissue clusters. (2) PIE visualized sample-wise functional enrichment, showing that sample points with simultaneously enriched functions were clustered together in the embedding space. This indicated the reasonableness of the PIE annotation and provided insights into the relationships between the functions.

**Figure 2.**
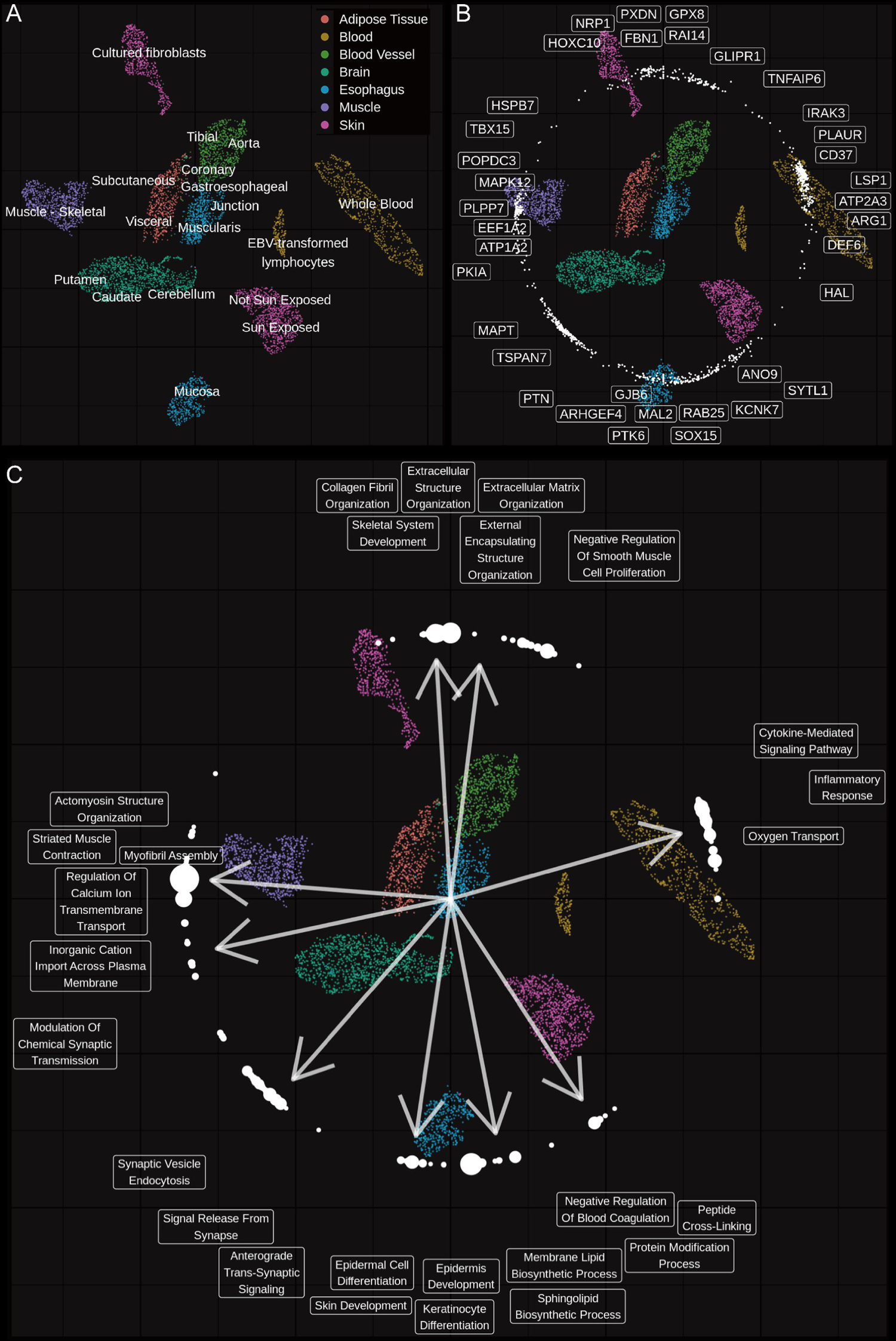
Functional Map of the GTEx embedding. A. 2D UMAP embedding of GTEx data. The processed GTEx dataset contained seven tissues that were further categorized into 27 tissue subtypes. Colors indicate different tissue types. As tissue subtypes existed, some tissues (Blood, Skin, Esophagus) had multiple distinct clusters. Subtype labels are shown near the corresponding samples in the embedding space. B. Overview of the mapped genes. In total, 908 genes were filtered using PIE for the GTEx data. In the embedding space, they are represented as points on a circle. The coordinates are the corresponding normalized weighting vectors. For visualization, we divided the circle into 12 equally spaced intervals based on the angles, selected the top three genes from each interval that had the highest association with the embedding structure, and displayed their names near their mapped coordinates. C. Overview of the functional map. Among the 5407 Gene Ontology Biological Process (GOBP) terms, 406 were found to be significantly enriched (p < 0.05) and are represented as points on the circle. The coordinates of the points are the normalized weighting vectors of the eigengenes of the corresponding functions. The sizes of the points correspond to the significance of the corresponding p-values for the gene sets. The larger the size, the smaller the p-values of the terms. From these 12 intervals, we identified the top three representative terms with the smallest p-values. In some intervals (e.g., the upper left), we did not find any enriched functions. In total, 27 representative terms were the 9 intervals. The arrows indicate the average mapped directions of representative terms for each interval.

PIE filtered and mapped 908 informative genes as dots on a circle in the embedding space. We identified genes with high Pearson correlations near their mapped coordinates (**Figure 2B**). Notably, some tissue-specific genes were mapped towards their corresponding tissues. For example, POPDC3, PLPP7, MAPK12, and ATP1A2 were mapped towards the muscle, whereas SOX15, MAL2, and RAB25 were mapped towards the esophagus (**see the human proteome atlas**, https://www.proteinatlas.org)^26^. After statistical testing, 406 of these 5407 GO terms were significantly enriched (p < 0.05, ratio = 7.51%). The mapped coordinates of 406 eigengenes are shown as dots in the circle in Figure 2C. To better visualize the representative enriched terms, we divided the circle into 12 equally spaced intervals based on angles. For each interval, we selected the three most significant terms with the lowest p-values. From these 12 intervals, we identified 27 representative enriched functions from nine intervals, with no significantly enriched functions identified in the remaining three intervals (**Figure 2C**).

An overview of the functional map suggested that the enriched functional directions highlighted the differences between the various clusters (**Figure 2C**). For instance, muscle tissue on the left was annotated with terms such as *Striated Muscle Contraction* (GO:0014888), *Myofibril Assembly* (GO:0030239), *Actomyosin Structure Organization* (GO:0031032), implying that muscle contraction is a key functional distinction between muscles and other tissues^27^. Conversely, the blood clusters, including EBV-transformed lymphocytes and whole blood, were annotated with functions related to circulation (*Oxygen Transport*, GO:0015671) and immunity (*Cytokine-Mediated Signaling Pathway*, GO:0019221; *Inflammatory Response*, GO:0006954), highlighting the crucial role of blood in transportation^28^ and the immune defense^29^. These annotated terms underscored the distinct functional differences between the clusters in their respective directions.

We visualized the sample-wise functional enrichments to focus and visualize individual functions. PIE calculates eigengene values^24^ as a quantitative measure of the functional enrichment for each gene set. From the 27 representative GO terms shown in Figure 2C, we selected four tissue-specific (**Figure 3A**) and four multi-tissue-specific terms (**Figure 3B**). Sample point colors represent functional enrichment levels, with brighter points indicating higher expression levels. We observed that terms such as *Striated Muscle Contraction* (GO:0014888) and *Oxygen Transport* (GO:0015671) were highly tissue-specific to muscle and blood, respectively. Additionally, the *Regulation Of Calcium Ion Transmembrane Transport* (GO:0070588) was enriched in both the muscle and brain, suggesting that similar calcium ion-regulated physiological activities, for instance, neuronal activities in the brain^30^ and muscle contractions^31^, might account for the proximities of these tissue clusters in the embedding space. Another example is the enrichment of *Negative Regulation Of Smooth Muscle Cell Proliferation* (GO:0048662) in blood vessels and esophageal muscularis, both components of the walls of blood vessels and esophageal muscularis layers^32,33^. Sample points with simultaneously enriched functions were clustered together in the embedding space, reinforcing that the PIE annotations are reasonable.

**Figure 3.**
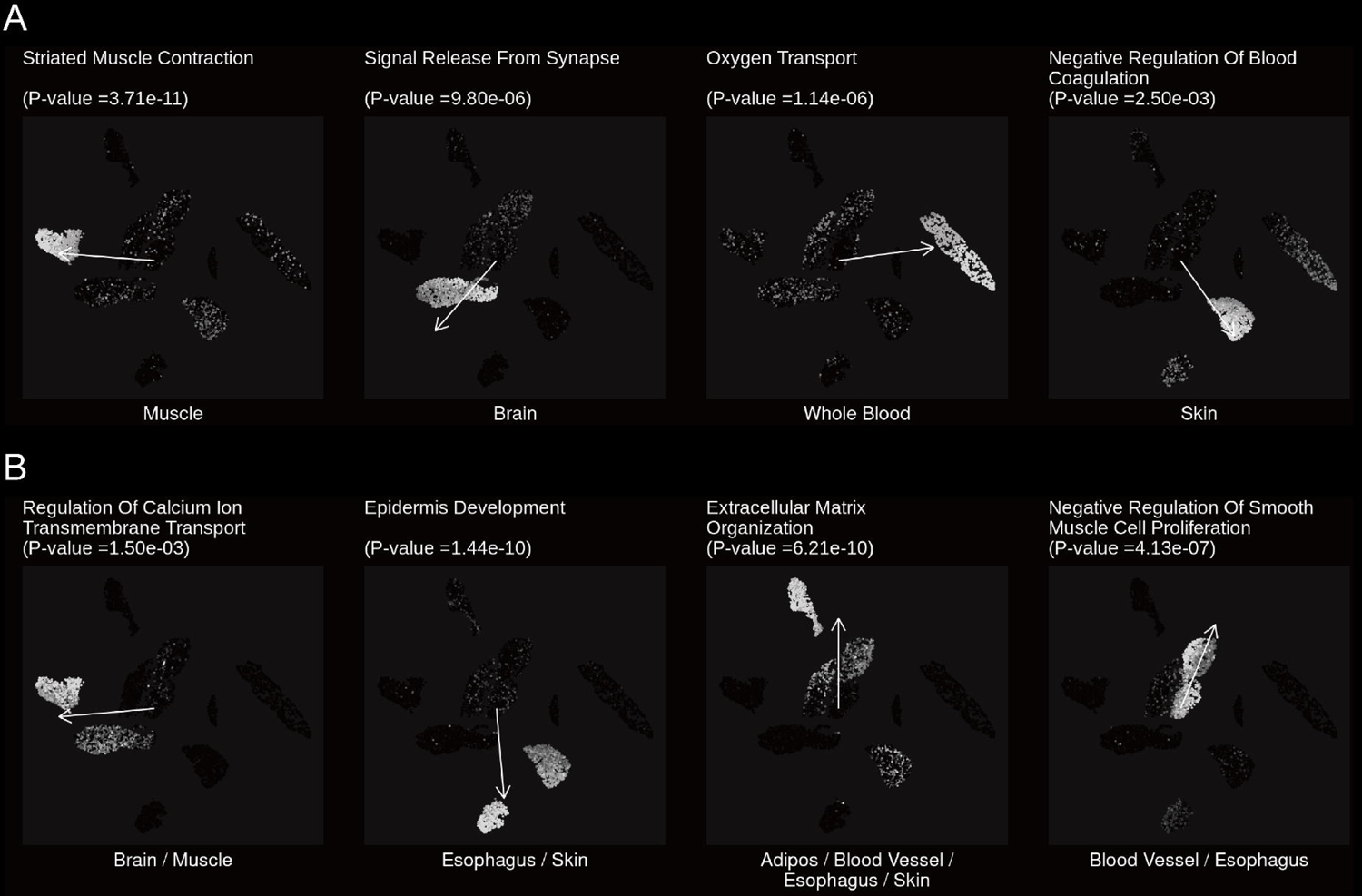
Visualization of the functional enrichment. A. Visualization of the four tissue-specific enriched functions. The colors indicate the eigengene values of the samples, that is, the sample-wise expression of the functional gene sets. Therefore, we used it to represent relative functional enrichment. Brighter colored sample points indicate higher functional enrichment. B. Visualization of the four multi-tissue-specific enriched functions. These functions were enriched in multiple tissues. Tissue clusters sharing similar functional enrichment were also close to each other in the embedding space. This suggests that functional annotation can be used to interpret cluster relationships in the embedding space.

To further investigate the functional relationships, we visualized the functions using a heatmap (**Figure 4A**) and a correlation plot (**Figure 4B**). Each row in the heatmap represents the eigengenes of the functional gene sets. Notably, functions with similar expression patterns pointed in a similar direction in the embedding space, as seen in Figure 2C. Moreover, the correlation plot of eigengenes suggested the presence of functional modules that were simultaneously enriched across the samples. This mapping by PIE between the biological functional space and the embedding space facilitates a deeper understanding of the relationships between functions.

**Figure 4.**
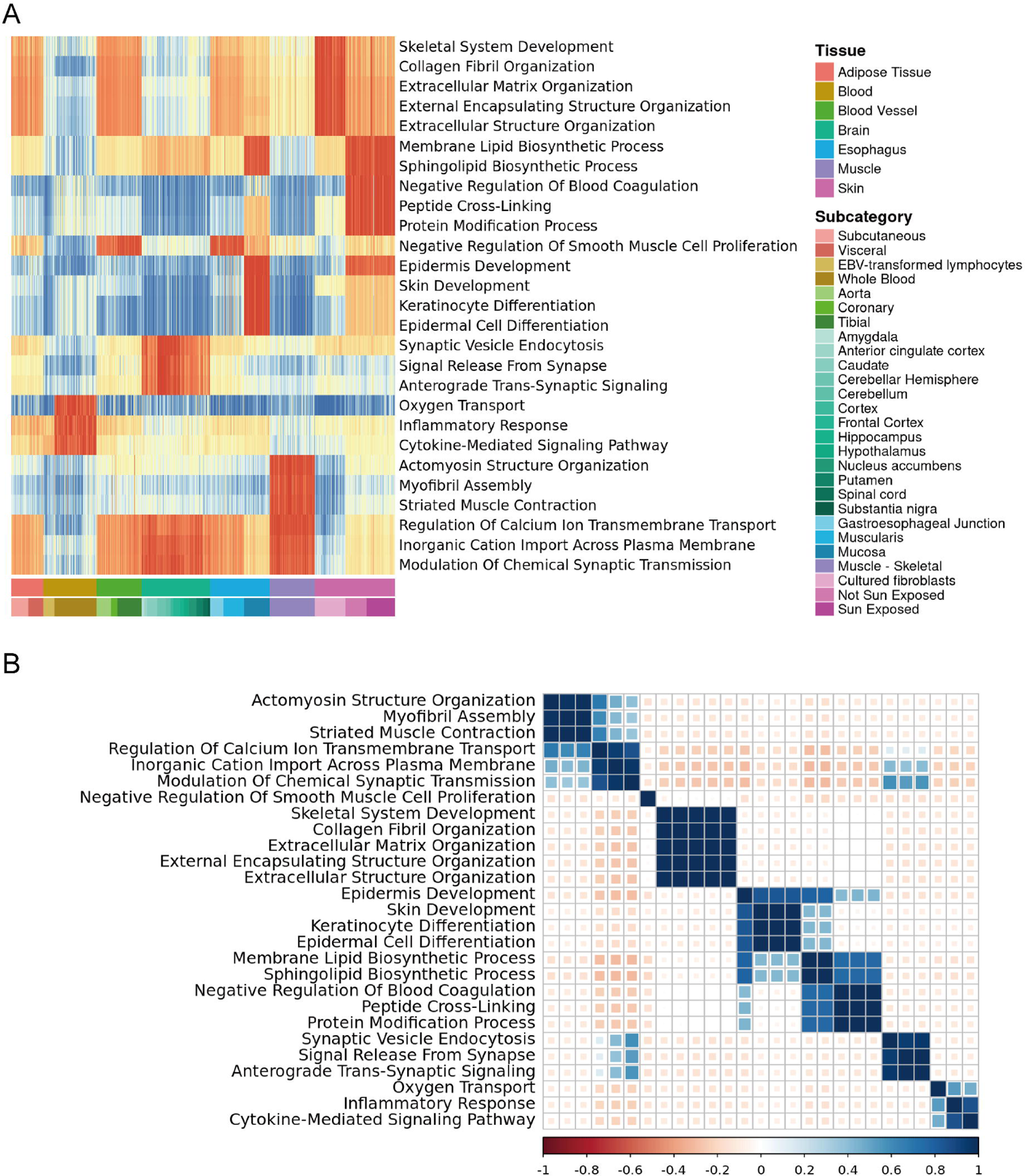
Overview of the top representative terms. A. Heatmap of the top terms. The heatmap was a 6220 × 27-sized matrix. Each row shows the eigenvalues of terms. The columns were ordered according to tissue subtype labels. Red/blue indicates relatively high/low expression levels of eigengenes across samples. B. Correlation plot of the top terms. Pearson correlation coefficients between the eigengenes were calculated and visualized as colors. Blue and red indicate a positive/negative correlation between the terms. The functional module structure shared similar expression patterns among samples.

### Annotation accuracy and reproducibility

Subsequently, the framework’s performance was evaluated by addressing two key questions: (1) Is the annotation of the PIE objective comparable to known knowledge? (2) Is the PIE robust and reproducible in interpreting embedding?

To evaluate PIE annotation, we used known tissue-specific genes from the Human Tissue-Specific Proteome (<u>HTSP</u>) in the Human Protein Atlas^26,34^. HTSP offers a comprehensive, objective definition of tissue-specific genes based on the transcriptome analysis of major organs and tissue types. Among the seven tissues, HTSP identified tissue-specific genes in the brain, muscle, esophagus, skin, and adipose tissues (**see method details**). For the 908 filtered informative genes of PIE, over 65% of the selected genes matched the HTSP-defined tissue-specific genes (**Figure 5A**). Fisher’s exact test confirmed that this selection was statistically significant compared to random guessing for most tissues (**Figure 5B**). The only exception was adipose tissue, which showed less separable in the embedding space compared to other tissues (**Figure 2A**), suggesting lower tissue-specificity and potentially complicating the identification of corresponding genes.

**Figure 5.**
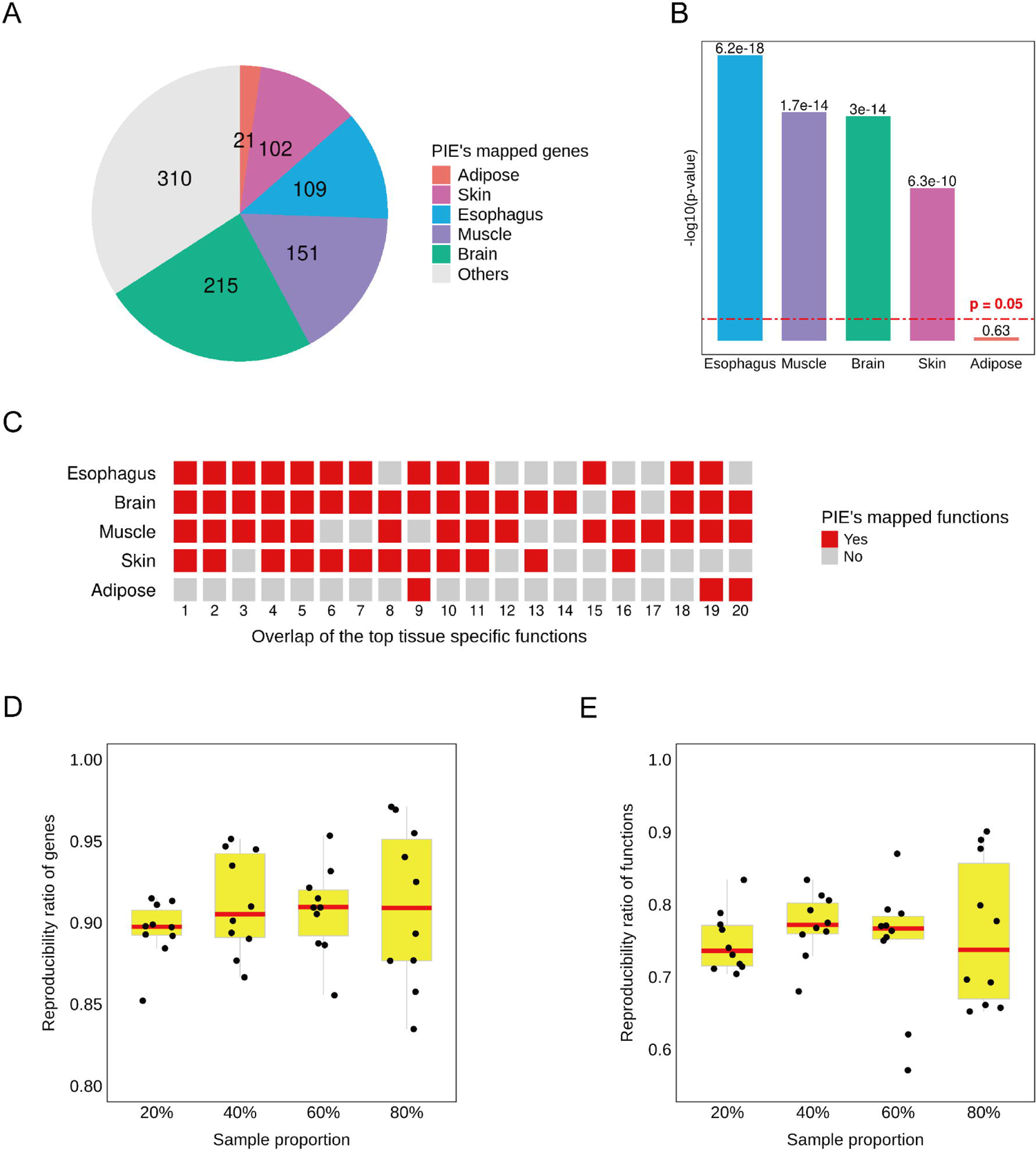
Evaluation for PIE on the GTEx data. A. Comparison of the selected genes. We compared the genes selected by PIE with tissue-specific genes defined using the human tissue-specific proteome (<u>HTSP</u>). Of the 908 genes, 598 matched the tissue-specific genes of HTSP in the brain, muscle, esophagus, skin, and adipose tissues. B. Fisher’s exact test for tissue-specific gene selection. From the 4549 genes in the GTEx data, 908 genes. We performed Fisher’s exact test to determine whether the selection strategy was effective in identifying tissue-specific genes for the brain, muscle, esophagus, skin, and adipose tissue. Except for adipose tissue, which showed lower specificity in embedding (Figure 2A), the gene selection strategy of PIE was effective. C. Comparison of the enriched functions. We compared the enriched functions identified by PIE and the tissue-specific functions defined based on the tissue-specific genes of HTSP. When comparing each tissue, we found that their functions matched well. The overlaps are denoted as red blocks; otherwise, they are denoted as gray blocks. Except for adipose tissue, most of the top 20 tissue-specific functions were identified using PIE. This suggests that PIE can be used to characterize tissue-specificity. D-E. Reproducibility of the selected genes and enriched functions. We randomly sampled and applied PIE to a subset of the GTEx data (sample ratio = 20%,40%, 60%, and 80%). We compared the reproducibility rates of the selected genes and enriched functions between the results of the subsets and those using all samples. To eliminate the randomness introduced by the sampling, UMAP, and permutation tests, we repeated them 10 times per sample ratio and showed the boxplots. For all sample ratios, the reproducibility of the genes and functions was above 85% and 70%, respectively.

To examine the enriched functions identified by PIE, we used HTSP tissue-specific genes for each tissue as input for Enrichr^35^ and identified the top 20 enriched functions with the smallest p-values. Consequently, we compared these overlapping functions to those identified and mapped by PIE in the embedding space. Except for adipose tissue, most of the top enriched functions were successfully identified by PIE (**Figure 5C**). Notably, PIE identified 18 of the top 20 brain tissue functions. Despite being unsupervised and requiring no tissue labels, PIE effectively identified tissue-specific functions that characterized distinct tissue clusters.

Finally, we evaluated the robustness and reproducibility of the proposed computational framework. We randomly sampled subsets from all samples, applied UMAP to obtain new embeddings, and performed PIE to identify enriched functions. To eliminate the effects of randomness and ensure a fair evaluation, we repeated this procedure 10 times for each sample size (sample ratio = 20%, 40%, 60%, and 80%). We compared the results from the randomly sampled subset with those from the entire sample. We calculated the reproducibility ratios for the filtered genes (**Figure 5D**) and the enriched functions (**Figure 5E**). The results showed that, even with a small sample proportion, the reproduced percentages of genes and their functions averaged above 85% and 70%, respectively.

### PIE interprets the developmental trajectories in the *C. elegans* embedding

For the second problem, we used a subset of the *C. elegans* embryogenesis dataset^18^, a single-cell RNA-seq dataset profiling ciliated neurons. The UMAP embedding effectively captured the complex developmental trajectories of the differentiated neuronal cells (**Figure 6A**). |n this analysis, we demonstrated that (1) PIE could map functions to interpret continuous developmental trajectories. (2) A limitation of PIE is its tendency to map only a single direction for each function, overlooking non-principal information within the corresponding functional gene set. (3) PIE can be applied to local embedding structures. Focusing on local embedding was beneficial, enabling us to extract informative genes and map relevant functions specific to the local structure. This approach allowed us to uncover and utilize non-principal information, often ignored in global embeddings, to enhance our understanding of local embeddings.

**Figure 6.**
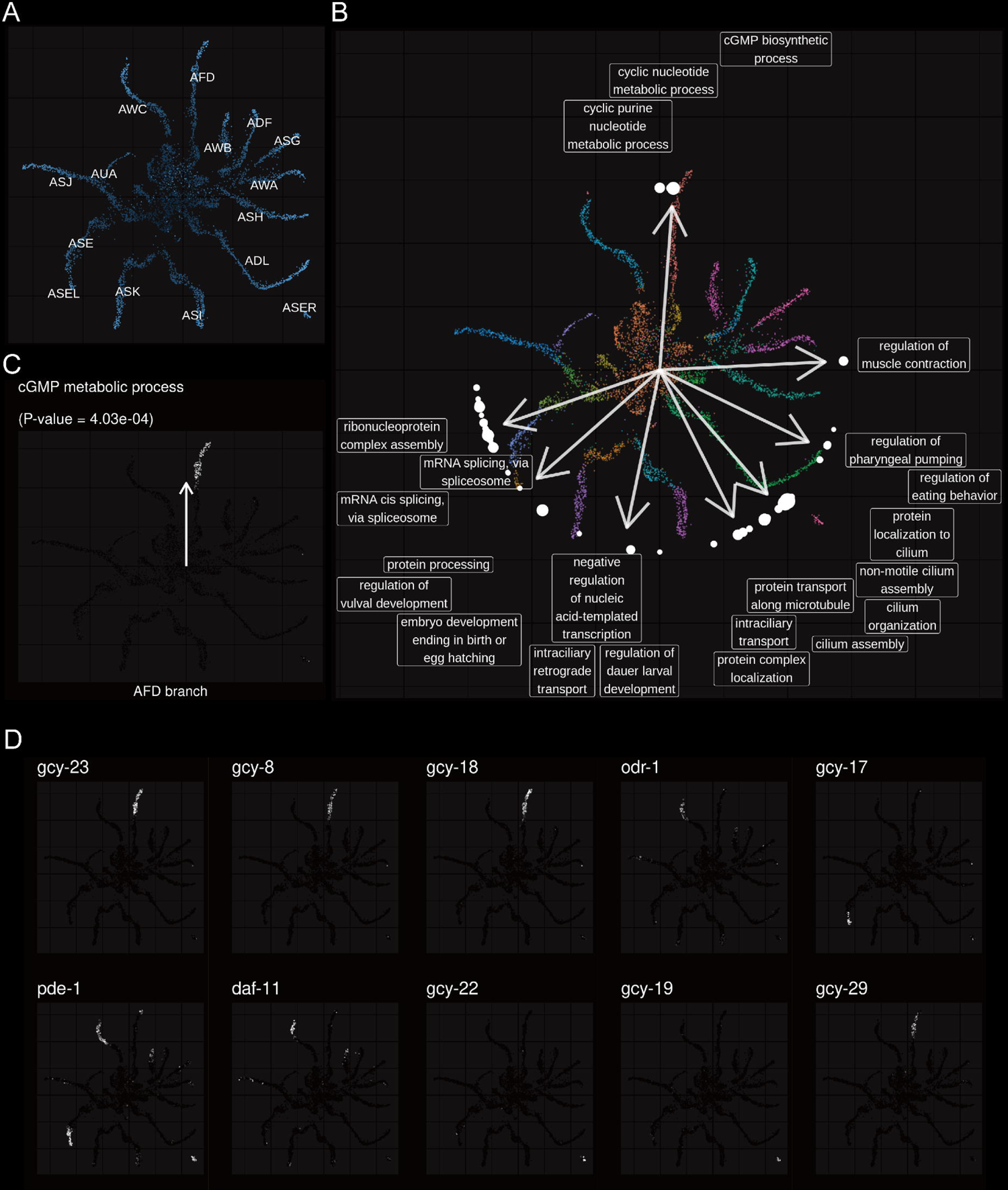
Functional Map of the *C. elegans* embedding. A. 2D UMAP embedding of the *C. elegans* data. The dataset includes 27 annotated neuronal cell types, some of which are shown in the embedding. The colors of the sample points indicate the embryo times, and brighter colors indicate that the samples were in the late developmental stage. B. Functional map of *C. elegans* embedding. Among the 1711 Gene Ontology Biological Process (GOBP) terms, 57 were significantly enriched (p < 0.05, ratio = 3.33%). White points on the circle represent the mapped coordinates of enriched terms. The sizes of the dots imply significance, and larger dots correspond to terms with smaller p-values. The 22 top representative terms from the eight intervals are shown. Arrows indicate the average coordinates of the top representative terms within each interval. C. Directions of *cGMP metabolic process* and functional enrichment. The *cGMP metabolic process* function (p = 4.03e-4) implied high enrichment in the AFD cell branch samples. D. Functional gene sets for the *cGMP metabolic process*. gcy-23, gcy-8, gcy-18 and gcy 29 showed similar high expression in the samples of the AFD, whereas odr-1, gcy-17, and pde-1 showed distinct expression patterns.

We first applied PIE to interpret the global structure of *C. elegans* embedding (**Figure 6A**). In this analysis, the sample points’ colors indicate embryo time and the trajectories of different neuron types aligned with the chronological order of embryo development. PIE filtered 861 informative genes from the original 8692 genes (9.91%) and identified 57 of the 1711 GO terms as significantly enriched (p < 0.05; ratio = 3.33%). These enriched terms are detailed across 12 intervals (**Figure 6B**). Among the enriched terms, we focused on the *cGMP metabolic process* (GO:0046068), which pointed to the AFD branch (**Figure 6C**). We investigated the overlapping informative genes within this functional gene set (**Figure 6D**). Previous studies have established that AFD, a key thermosensory neuron, relies on cGMP signaling for thermosensation^36^. PIE confirmed that AFD cells showed high functional enrichment of the *cGMP metabolic process* compared to other cells. Further analysis of gene expression patterns found that gcy-8, gcy-18, gcy-23, and gcy-29 were highly expressed in AFD cells. The gcy-8, gcy-18, and gcy-23 genes are crucial for thermotaxis in *C. elegans*^37^.

Contrastingly, genes showing different expression patterns were highly expressed in other cells (**Figure 6D**). These genes (e.g., odr-1, pde-1, and daf-11) were overlooked when mapping the enriched functional directions. As PIE leverages the eigengene, the first principal component score was used to summarize the functional gene set. Consequently, some genes were ignored despite their potential association with function.

Finally, the PIE was applied to the local embedding structure. The neurons on the left-bottom of the *C. elegans* embedding were selected to gain a “zoom in” view (**Figure 7A**). PIE application found 66 enriched representations of 22 terms (**Figure 7A**). Interestingly, terms such as *cilium assembly* (GO:0060271) were enriched in both local (**Figure 7B**) and previous global embedding (**Figure 6A**) but highlighted different trajectories. However, annotation for the local structure could be obtained by focusing on a local structure.

**Figure 7.**
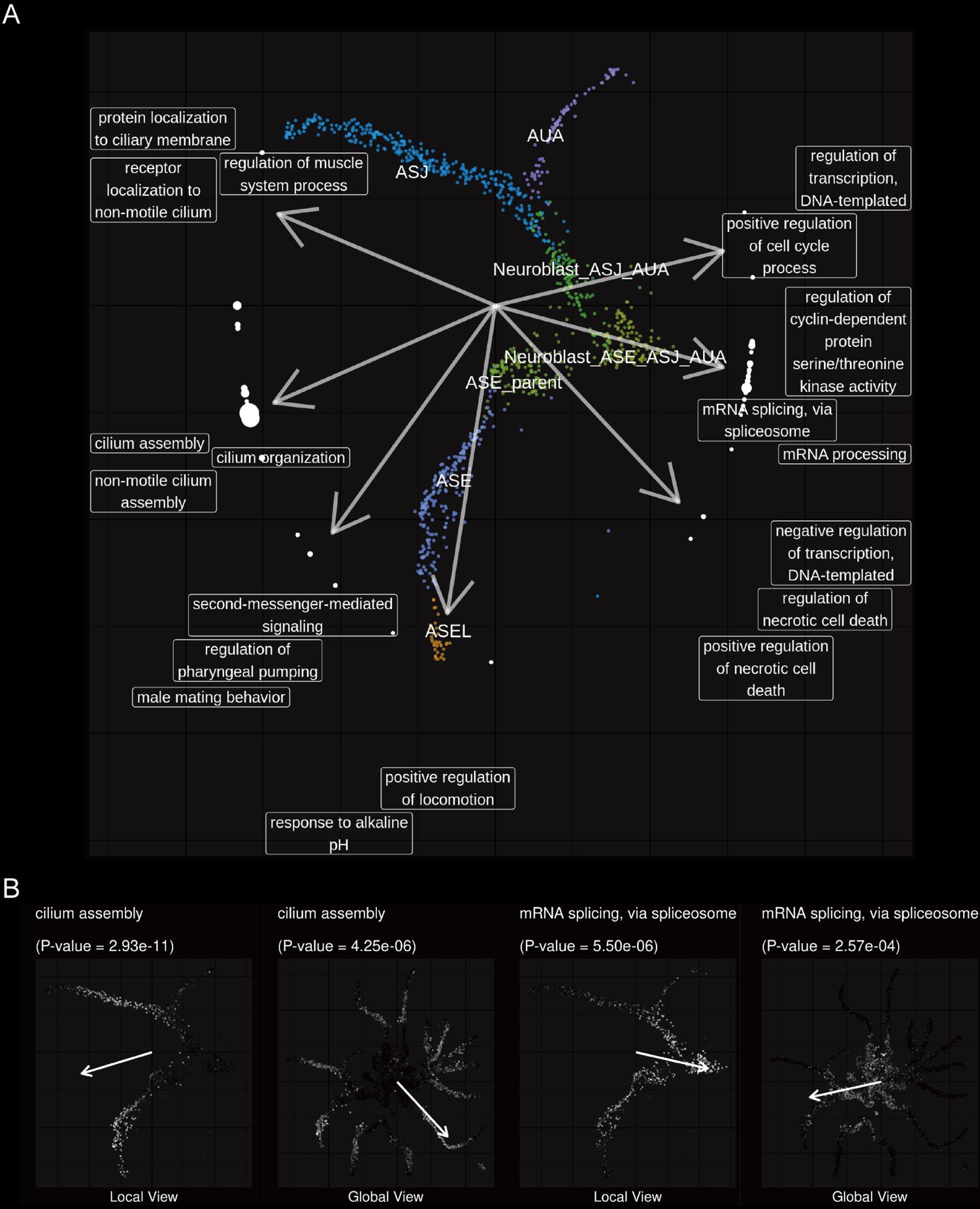
Functional Map of the local embedding. A. Overview of functional map of local embedding. The local structure focuses on the seven neurons at the bottom left of the original global embedding. The colors of the dots showed the 7 different cell types. In total, 66 GO terms were significantly enriched (p < 0.05), and 19 representative terms are shown. B. Comparison of enriched directions of functions for local and global embedding. The terms *cilium assembly* and *mRNA splicing, via spliceosome* were enriched in local and global embedding, but their mapped enriched directions were different. This suggested that the “zoom in” view helped obtain insight into the local structure.

## Discussion

In this study, we proposed and demonstrated a computational framework to interpret nonlinear DR embeddings. Despite the widespread use of DR techniques, interpreting the structure of nonlinear DR embedding remains challenging, preventing us from completely understanding the data. There is a notable knowledge gap between the low-dimensional embedding space and the associated biological functions. To address this, we developed PIE, which provides functional annotations of the embedding space and easy-to-understand visualizations to elucidate the embedding structure. To our knowledge, this is the first methodological framework to provide post-hoc interpretations for embeddings through systematic functional mapping.

The interpretations provided by PIE benefit users in exploratory data analysis. We demonstrated the effectiveness of our method in the GTEx and *C. elegans* datasets. In both biological settings, PIE provided meaningful annotations for the embedding spaces, undercovering the functional differences between distinct clusters and trajectories. PIE identifies relatively enriched functional directions, enabling the users to characterize the hidden structure from a systems biology perspective. Exploring the connections between biological functions and their representations is important in systems biology. In this study, we focused on the embedding of sample points as the biological representation. In a different context, SAFE^38^ addressed the functional annotation of biological networks, such as gene–gene and protein-protein. The motivations behind PIE and SAFE were remarkably similar. While PIE interprets the functional differences in sample structure, SAFE interprets the functional attributes of local structure in gene and protein networks. Both approaches aim to enhance our understanding of biological data through functional annotations.

Conventional frameworks for comparing functional differences between groups involve two steps: first, identifying differentially expressed genes or proteins between the two groups (e.g., through controlled experiments); second, using the gene or protein list to perform statistical tests to identify differentially enriched GO terms and pathways. This approach has limitations, as it overlooks within-class differences between samples and requires label information. Moreover, it becomes complicated when comparing multiple groups. PIE offers a unique approach: it is unsupervised, requires no label information, and it provides a global functional map for any complex dataset rather than being restricted to two-group comparisons. Unlike conventional methods, PIE explores functional differences based on the embedded structure of the data, offering insights into the underlying structures.

Moreover, we envision several improvements to the proposed approach. First, the computational framework could be made interactive. For instance, we showed that the “zoom in” enabled a local interpretation of the embedding (e.g., a single cluster structure or a lineage of trajectory). However, as noted, PIE currently maps each function in one direction within the embedding space. It might be reasonable to consider mapping certain important functions in multiple directions. Additionally, a deeper understanding of the functional and embedding spaces would enhance the framework, allowing for the replacement and improvement of components such as feature selection, feature mapping, and statistical testing steps.

## Acknowledgments

This work was supported by the Human Glycome Atlas Project (HGA) and JSPS KAKENHI, 442 Grant Number: JP20H04282.

## Author contributions

Conceptualization, B.Z., K.U., and Y.M.; Methodology, Software, Formal Analysis, Data Curation, Writing – Original Draft, B.Z.; Investigation, B.Z. and K.H.; Visualization, B.Z., H.K. and Y.M.; Writing – Review & Editing, B.Z., K.U., H.K., K.H. and Y.M.; Supervision, Resources, Project Administration, Funding Acquisition, Y.M.

## Declaration of interests

The authors declare no competing interests.

## Method details

The computational framework required three key inputs: an expression data matrix, an embedding matrix of sample coordinates, and functional gene sets. The framework involved two main steps: (a) PIE selects and maps the genes to the embedding space and (b) PIE identifies the enriched functions and visualizes them in the embedding space. The input data and detailed computational procedures are described as follows:

### Input data

1. An expression matrix *X* ∈ *R*^*n*×*p*^ consists of *n* data points and *p* features. These features can be attributed to either genes or proteins. We denote the matrix by bold uppercase *X*, with *x*_.*l*_ denoting its *l*-th column vector and *x*_*m*._ denoting its *m*-th row vector. The element at position (*i*, *j*) in *X* is denoted by *x*_*ij*_.
2. An embedding matrix *Y* ∈ *R*^*n*×*d*^ containing d-dimensional space. The matrix represented the low-dimensional coordinates of the samples, obtained using the nonlinear DR method.
3. Functional gene sets in the form of a matrix *A* ∈ {0,1}^*F*×*G*^. This binary matrix represents the relationship between F functions and *G* genes. If the f-th function is associated with the *g*-th gene, then *A*_*f*×*g*_=1, otherwise, *A*_*f*×*g*_ = 0. These gene sets can be user-defined or sourced from existing databases.

### Filtering and mapping genes

To filter the informative genes, SCMarker^21^ was applied to the expression matrix *X* ∈ *R*^*n*×*p*^. SCMarker is an unsupervised feature filter that was initially designed for single-cell RNA-seq data. It identifies discriminative markers having distinctive expression patterns across samples, particularly markers that are co- or mutually exclusively expressed. SCMarker accurately identified features that maintained the embedding structure in the experiment. *X*^∗^ ∈ *R*^*n*×*p*∗^ denoted the filtered expression matrix consisting of *p*_∗_ informative genes.

Projection pursuit was applied to map the genes^23^. For two matrices *X* ∈ *R*^*n*×*p*1^, *Y* ∈ *R*^*n*×*p*2^, projection pursuit found weighting vectors *a* ∈ *R*^*p*1^, *b* ∈ *R*^*p*2^ such that the linear combinations *Xa*^*t*^ and *Yb*^*t*^ have a maximum association *C*(*Xa*^*t*^, *Yb*^*t*^). The association measure *C* can be the Pearson, Spearman, or Kendall rank correlation. Projection pursuit considers the following optimization problem:

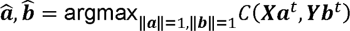

We consider a special case and compare each univariate gene vector, *x*^∗^ of *X*^∗^ and the embedding matrix *Y*. Therefore, we find the ^*b*^^_*l*_, *l* = 1, …, *p*_∗_:

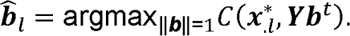

When the association measure is Pearson’s correlation, the normalized weighting vector ^*b*^^_*l*_ can be viewed as the normalized coefficient vector of the linear regression model. ^*b*^^_*l*_ provides a measure of how embedding describes a gene vector. Here, we take the ^*b*^^_*l*_ as the representation of the l-th gene. This idea can be analogized to using loadings as representations of features in a PCA biplot^41^.

Meanwhile, genes whose weighting vectors ^*b*^^ have a higher association *C* (*x*^∗^, *Y*^*b*^ *t*^) imply high consistency with the embedding, hence, are more important. Genes with weighting vectors implying higher association measures were filtered out. Clearly, ^*b*^^_*l*_ is uniformly constricted, where in a 2D or 3D embedding space, ^*b*^^_*l*_ is always located on a unit circle or ball.

### Performing enrichment analysis of functional gene sets

Functional gene sets were mapped to an embedding space, thus filtering out informative genes. For the functional analysis, we identified informative genes in the corresponding functional gene set. To characterize the high view of the function, we leverage the concept of an eigengene. Let the submatrix of the *f*-th gene set be *X*_*f*_ ∈ *R*^*n*×*pf*^, where *n* is the sample size and *p*_*f*_ is the size of the informative genes in the functional gene set. The singular-value decomposition of *X*_*f*_ is given by

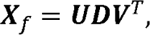

where *U* is a matrix with orthonormal columns *U* = [*u*_1_, …, *u*_p*f*_], and *D* = diag{*d*_1_, *d*_2_, …, *d*_*pf*_ } is a diagonal matrix in which *d*_1_ > *d*_2_ > ⋯ > *d*_*pf*_ are singular values in decreasing order. *V* is an orthogonal matrix. We used *u*_1_, the eigengene, to characterize and represent the functions.

Furthermore, the eigengenes were mapped to the embedding space. The process was similar to mapping of single genes using Projection Pursuit. Given the weighting vector ^*b*^^_*f*_ of the *f*-th eigengene, we measured the distance between ^*b*^^_*f*_ and ^*b*^^_*i*_, *i* = 1, …, *p*^∗^, where the *p*^∗^ indicated the number of informative genes. We used the cosine similarities *d*_f,i_ = cos(^*b*^^, ^*b*^^) = ^*b*^ *t*^ ^*b*^^. We then defined a gene rank based on this distance.

For each functional gene set, after obtaining the gene rank, we perform a permutation test of the Gene Set Enrichment Analysis (GSEA) to identify whether the corresponding functions are significantly enriched (p-value < 0.05) in their mapped directions.

### Visualization of functions

Data visualization enhances the user’s ability to gain insights and understand data intuitively. We used two visualization techniques for representing functions in the embedding space: arrows and dot plots. For an enriched function, we used the mapped coordinates of its eigengene as the end of an arrow, with the origin as the starting point. We highlighted the highly expressed sample points in the embedding space by assigning them bright colors, indicating relatively high eigengene expression. The arrows indicate the direction of functional enrichment, while the colors indicate the sample-wise degree of functional enrichment.

To visualize the functional map in the embedding, we divided the entire space into 12 equally spaced intervals based on angles (30 °per interval). For each interval, we selected the top three representative terms with the smallest p-values from those whose enriched directions fell within the interval. This method was used to generate the visualizations in Figures 2A, 6A, and 7A.

### Implementation of PIE

We used UMAP to obtain a 2D embedding of the dataset, utilizing the umap function from the uwot R package. We reduced the dimensionality to a 2D embedding space (n_components = 2), set the neighbor number for the k-nearest neighbor graph to 15 (n_neighbors = 15), and the effective minimum distance to 0.5 (min_dist = 0.5), while keeping other parameters at their default. In the filter step, we used the SCMarker R package with the following parameters: geneK = 10, cellK = 10, maximum number of nearest neighbors (K = 300), and minimum number of coexpressed/mutually exclusive cells (n = 10). For the optimization of projection pursuit, we used the ccaPP R package^23^, and used the Pearson correlation as the association measure (method = “person”). We used the fgsea R package^42^ to perform the GSEA test, restricting the size of the gene set to between 3–300 (minSize = 3, maxSize = 300), excluding any sets outside this range. The p-value threshold was set to 0.05, and identifying genes with p-values below this threshold was significantly enriched. PIE was applied to all the datasets with the same parameter settings. All associated R codes for producing all Figures are available on https://github.com/bingyuan-zhang/pie.

### Dataset processing

The GTEx dataset is a bulk RNA-seq dataset consisting of transcripts per million (TPM) from the original GTEx V8^17^. We focused on the protein-coding genes by utilizing the *protein-coding gene* list from the HGNC data based^43^, which defined by HUGO Gene Nomenclature Committee (https://www.genenames.org/download/statistics-and-files/). To ensure sample quality, we selected samples with RNA integrity numbers (*RIN* > 7). We focused on major tissue types by excluding tissues with fewer than 500 sample points. To reduce noise, we filtered out genes with low variance (bottom 25%). The procedure retained 6220 samples and 4549 genes from 7 tissues, encompassing 27 subcategories.

The *C. elegans* dataset comprised single-cell RNA-seq count data, specifically a subset of the original *C. elegans* dataset^18^ as examined in Monocle 3^44^. We removed genes with total counts across samples less than 10 and removed genes with low variances (bottom 25%) to reduce noise. This procedure retained 6188 samples and 8692 genes. A batch correction was performed following the same procedure as outlined in the Monocle 3^45^ example (https://cole-trapnell-lab.github.io/monocle3/docs/trajectories/).

### Functional gene sets

In practice, functional gene sets can be defined by the user based on the relevant functions of interest or sourced from public knowledge bases such as Gene Ontology^19^, KEGG^39^ and Reactome^40^.

For the GTEx data, we used *GO_Biological_Process_2023* (https://maayanlab.cloud/Enrichr) sourced from Enrichr^35,46^, which defines 5407 functional gene sets for humans. For the *C. elegans* data, we used the *GO_Biological_Process_2018* (https://maayanlab.cloud/WormEnrichr/) dataset from *WormEnrichr* ^35,46^, which defines 1711 functional gene sets for Caenorhabditis elegans.

### Definition of the tissue-specific genes

Tissue-specific genes were identified using the Human Tissue-Specific Proteome (<u>HTSP</u>) from the Human Protein Atlas^26,34^. The HTSP classifies 20,162 human genes based on their expression patterns across 37 major organs and tissue types. It defines three tissue-specific subcategories based on their elevated expression levels: tissue enrichment, group enrichment, and tissue enhancement. We considered gene A to be specific to tissue B - if it fell into one of the HTSP-defined subcategories for tissue B.

## Notes

### Competing Interest Statement

The authors have declared no competing interest.

